# Immunoinformatic approach to design a vaccine against SARS-COV-2 membrane glycoprotein

**DOI:** 10.1101/2021.03.26.436314

**Authors:** Radhika Ravindran, Shoba Gunasekaran, Murugesh Easwaran, Sajitha Lulu, P. Ambili Unni, S. Vino, Mukesh Doble

## Abstract

SARS-COV-2 is a pandemic virus causing COVID-19 disease which affects lungs and upper respiratory tract leading to progressive increase in the death rate worldwide. Currently, there are more than 123 million cases and over 2.71 million confirmed death caused by this virus. In this study, by utilizing an immunoinformatic approach, multiepitope-based vaccine is designed from the membrane protein which plays a vital role in the virion assembly of the novel-CoV. A total of 19 MHC class- I binders with HLA-A and HLA-B alleles have been selected with NetMHC pan EL 4.0 method from IEDB MHC-I prediction server. Four epitopes candidates from M-protein were selected based on the antigenicity, stability, immunogenicity, Ramachandran plot and scores with 100 % was taken for docking analysis with alleles HLA-A (PDB ID: 1B0R) and HLA-B (PDB ID: 3C9N) using ClusPro server. Among the four epitopes, the epitope FVLAAVYRI has the least binding energy and forms electrostatic, hydrogen and hydrophobic interactions with HLA-A (−932.8 Kcal/mol) and HLA-B (−860.7 Kcal/mol) which induce the T-cell response. Each HLA-A and HLA-B complex in the system environment achieves stable backbone configuration between 45-100 ns of MD simulation. This study reports a potent antigenic and immunogenic profile of FVLAAVYRI epitope from M-protein and further *in vitro* and *in vivo* validation is needed for its adaptive use as vaccine against COVID-19.

## 1.0 INTRODUCTION

Coronaviruses (CoVs) are a group ofRNA viruses with genomic size ranging approximately between 26 to 32 kb. Based on the criteria of genetic and antigenetic property, CoV is organised into alpha and beta coronaviruses which infect mammals; gamma and delta coronaviruses which infect birds. The viral envelope is bilayered with outer Membrane glycoprotein (M), Envelope (E), Spike (S) protein and the inner Nucleocapsid (N) protein (Baker 2008). Each of the above four proteins is vital to maintain the structural integrity of viral cell membrane and also in the replication cycle. In this current work, we have focused on developing an epitope-based vaccine using the Membrane glycoprotein (M). M protein is a major transmembrane glycoprotein with multiple biological functions. It is present abundantly on the surface of SARS- COVID and plays an important role in viral assembly (Neuman et al. 2011). M protein is able to induce antigen response during infection or immunizations (Wesseling et al. 1993).Epitope/peptide based vaccines can effectively engage in inhibiting the viral replication and they are economical when compared to conventional vaccines. These types of vaccines can be processed very rapidly, are cheap and economical without undergoing any *invitro* culturing of a pathogenic SARS-CoV virus. The selectivity of the epitope/peptide requires the explicit immunological trigger responses through the selection of immunodominant and conserved epitopes (Skwarczynski and Toth 2016).The success of the prepared vaccine will depend on properties such as stability, immunogenicity and allergenicity (Oyarzun et al. 2015).

In this study, B cells and helper T cells analysis are used in the selection of epitopes for designing an *in-silico* vaccine. The selected epitopes should activate T cells potentially that are required to stimulate a protective immunity (Li et al. 2014). The principal action of the peptide vaccines is based on the synthesis of the determined B and T cell epitopes which are immunodominant and should produce specific immune responses. Epitopes from B cells (it can be proteins) can be combined to T-cell epitope (short peptide of 8-20 amino acids) to make the prepared vaccine immunogenic in nature (Dermime et al. 2004).

## 2.0 Materials and methods

The flow chart summarises the methods, tools and servers used for selecting the epitopes (**Fig.1**).

**Fig. 1.**
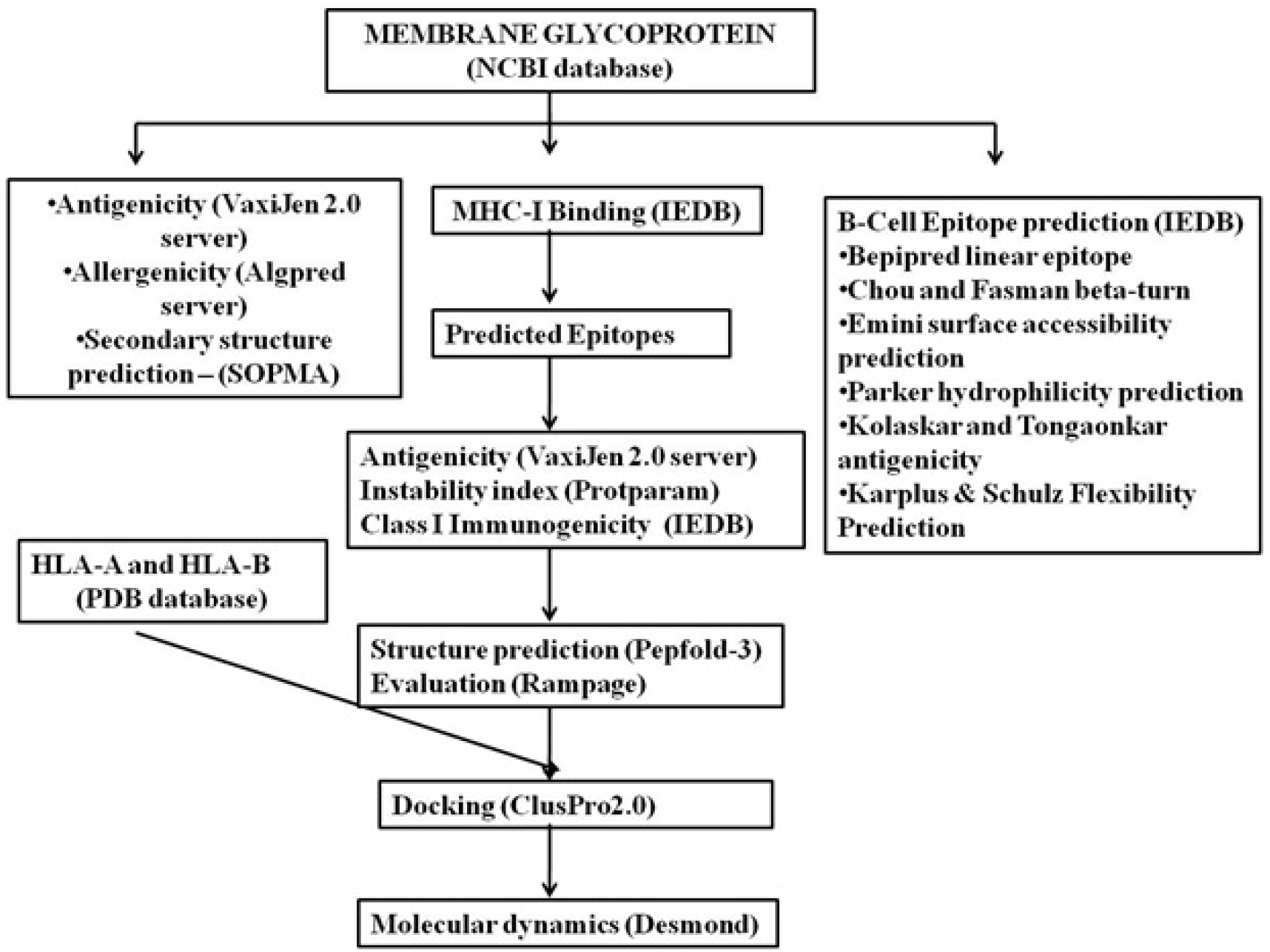
Flow chart of the methods, tools and servers used for epitope predictions.

### 2.1 Dataset

The sequence of membrane glycoprotein from Human SARS coronavirus (SARS-CoV) was retrieved from NCBI database and its Genbank ID was found to be QKR85720.1 (https://www.ncbi.nlm.nih.gov/protein/QKR85720.1). Membrane glycoprotein plays a significant role in the viral assembly and morphogenesis and is reported for its pathogenesis associated with the SARS-CoV infection. Protein sub cellular location was predicted using Virus mPLoc which accurately predicts viral proteins location in a host cell(Shen and Chou 2010).

### 2.2 Allergenicity and antigenicity prediction

Allergenicity prediction was performed by utilizing AlgPred server, which utilizes Support Vector Machine (SVM) and motif based methods for categorizing allergens and non-allergens (Saha and Raghava 2006). VaxiJen v 2.0 server is used for predicting the antigenicity of epitope, based on autocross covariance (ACC) transformation of the protein sequences into uniform vectors(Doytchinova and Flower 2007).

### 2.3 Secondary structure and stability prediction (SOPMA)

Secondary structure of the membrane protein was predicted using SOPMA.It predicts percentage of secondary structure elements such as alpha helix, beta sheet, coils and turns (Rozas and Rozas 1995). The instability index of membrane protein was predicted using ProtParam tool.

### 2.4 T- cell epitope prediction

The T-cell epitope represents smaller peptide sequences, which could bind with the Major Histocompatibility Complex (MHC) present on Antigen Presenting Cells (APC) to evoke the immune responses. The T cell epitope prediction was performed using IEDB server, which predicts the binding of epitopes to human leukocyte antigens HLA-A and HLA-B present on class I MHC (Atanasova et al. 2013). Epitopes with maximum number of binders in MHC class I alleles were considered for further analysis.

### 2.5 Modeling and validation of epitopes

The three dimensional structure of epitopes which are binders of class II MHC were predicted by using PEP-FOLD server (Lamiable et al. 2016).It aids in predicting the structure of linear and disulphide bonded cyclic peptides of length ranging between 9-36 amino acid (Thevenet et al. 2012).

The stereochemical properties of modelled epitopes were analysed by performing Ramachandran plot analysis. RAMPAGE server was employed for the retrieval of stereochemical features of the modelled epitopes (Lovell et al. 2003). This server also assesses the quality of the modelled epitopes by considering the frequency of amino acids falling in the favoured, allowed and disallowed regions of the Ramachandran plot.

### 2.6 Molecular Docking study

Molecular docking studies provide significant insights into the protein-ligand interactions and underlying biochemical mechanisms. The binding affinity between HLA alleles of class II MHC and epitopes were analysed by conducting molecular docking studies using a web based server ClusPro (Kozakov et al. 2017). Docking algorithm employed in ClusPro identifies the conformers with least energy and favourable surface complementarities. The molecular interaction between the predicted epitopes and HLA A (PDB ID: 1B0R) and HLA B (PDB ID: 3C9N) was performed.

### 2.7 B cell epitope prediction

The modules from IEDB were employed for predicting B cell epitope by Kolaskar and Tongaonkar module (Kolaskar and Tongaonkar 1990). Further, properties of the B cell epitopes such as surface accessibility, flexibility and hydrophilicity were predicted using Emini surface accessibility (Emini et al. 1985), Karplus and Schulz flexibility (Karplus and Schulz 1985) and Parker hydrophilicity prediction (Parker et al. 1986) respectively.

### 2.8 Molecular dynamics study

The structural stability of both, HLA-A and B epitopes and protein complexes were calculated by Desmond software, an embedded version from Schrodinger suite. Water solvent of 3 to 6 site configuration was used to build the system with electrostatic constant of 332.1A°. The geometry parameters were maintained to keep 0.95 A° for r(OH), 104.52 A° for HOH; constants for electric dipole moment were kept stable upto*-*0.80for q(O) and +0.40 for q(H). The trajectory samples were kept for 1000 frames; 100 nanosecond simulation time; recording trajectory interval for 100 pico second with 1.2 dynamic energy state for each trajectory. Isothermal-Isobaric (NPT) ensemble was selected to contact thermostat temperature of regular interval period. The system was kept to observe at 300 K with 1.013 bar pressure. The resultant structures were subjected to study displacement of complete protein complex, local changes along the protein complex chain, secondary structure elemental changes such as alpha-helical and beta strand movements, and the index of residual state in the displacements observed through RMSD and RMSF.

## 3.0 Results

### 3.1 Dataset

Human SARS coronavirus (SARS-CoV) membrane glycoprotein was considered for this study. The protein of SARS-CoV is retrieved from NCBI and its Gene bank accession number is QKR85720.1. It is a type III transmembrane glycoprotein with 222 long amino acidsand, they reside in the host cell membrane. The dimerization of M protein and its interaction with E protein plays a role in maintaining the spherical shape of the virus and the formation of a matrix-like layer underneath the viral membrane.(J Alsaadi and Jones 2019; Neuman et al. 2011).

### 3.2 Determination of protein antigenicity and allergenicity

The antigenicity and the allergenicity of SARS-CoV membrane protein was predicted by employing VaXiJen v 2.0 and AlgPred server respectively. The antigenic score of membrane glycoprotein is 0.5047. AlgPred server is based on SVM based algorithm for prediction of allergenicity and it was found to be non-allergen.

### 3.3 Secondary structure analysis of complete membrane protein and peptides predicted for vaccine candidate

The secondary structural features of SARS-CoV2 membrane glycoprotein was found to be 33.78 % of alpha helix, 22.97 % of beta sheet, 4.95 % of turn and 38.29 % of coil.The observation exhibits that the vaccine candidates are profiled in the geometry of helical and stable configuration which interact with the alleles HLA-A and HLA-B respectively.

### 3.4 T- cell epitope prediction

The selection of the predicted epitopes were accomplished by considering their affinity towards class I MHC. MHC binding T cell epitopes possessing scores closer to 1 have higher affinity and only those were considered for further studies (**Table 1**).The complete T cell epitope prediction were provided in the **Supplementary Table.**

**Table 1:**
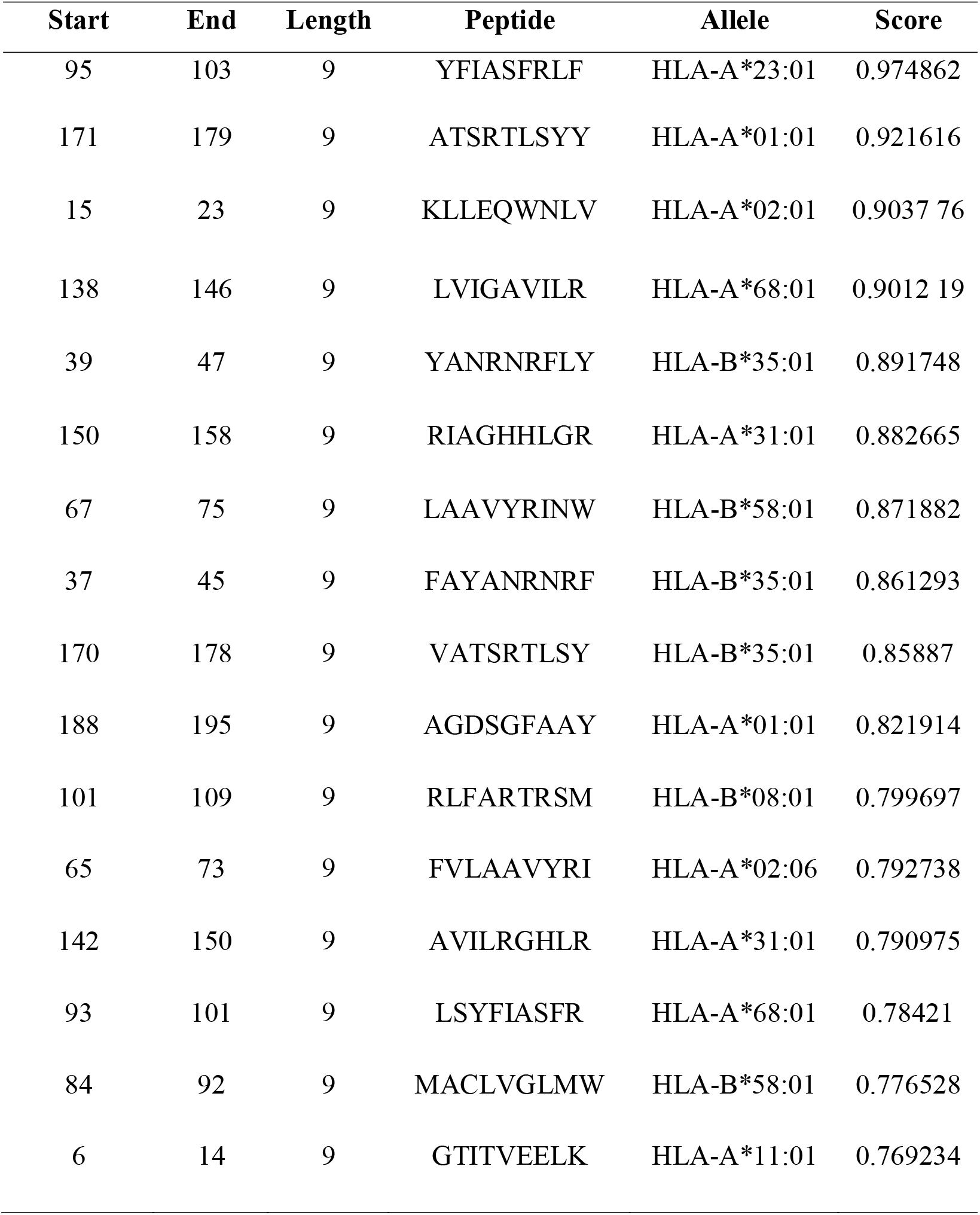
T cell epitopes predicted for SARS-CoV2-membrane glycoprotein.

### 3.5 Determination of antigenicity, instability index and immunogenicity of predicted epitopes

The antigenicity, instability index and immunogenicity of the best short listed epitopes were analysed using VaxiJen, Protparam and IEDB tools respectively. Vaxigen is a server used to predict antigenicity of given peptide and its score is greater than 0.4 and considered for further analysis. Protparam computes various physico – chemical properties between the peptide sequences and whose instability index is smaller than 40is predicted as stable. Immunogenicity score is a predictor of immunogenicity, higher the score, greater the immunogenicity in the predicted peptide (**Table 2**). Epitopes such as LAAVYRINW, AGDSGFAAY, FVLAAVYRI, LSYFIASFR, GTITVEELK, and GFAAYSRYR exhibited good antigenic and immunogenicity scores when compared to the other epitopes. Hence these epitopes were considered for further analysis based on antigenicity, stability and immunogenicity score. Among the above six epitopes, the four epitopes AGDSGFAAY, FVLAAVYRI, LSYFIASFR, and GFAAYSRYR has 100% of favored regions in the Ramachandran plot which are considered for further analysis.

**Table 2:**
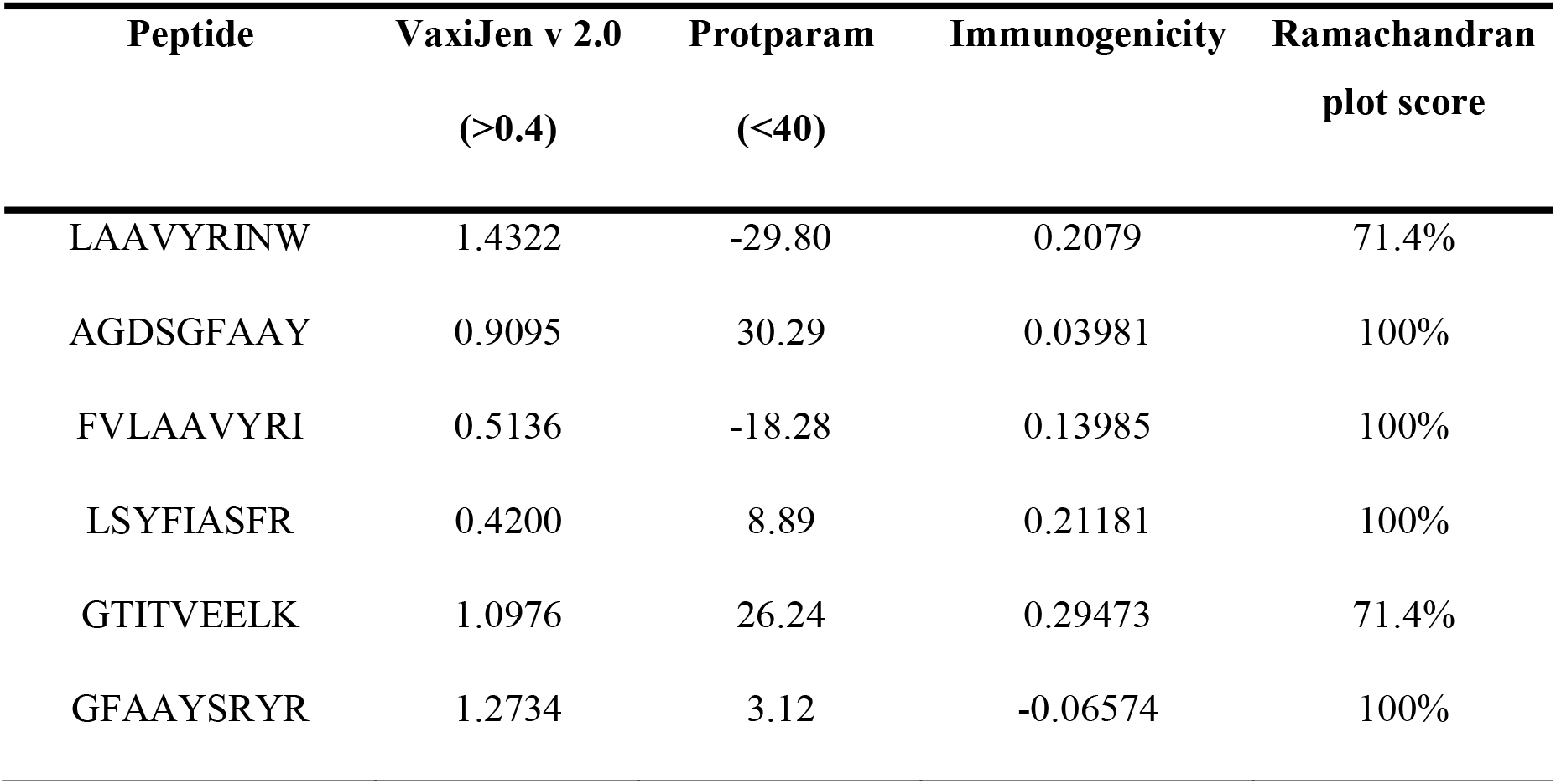
Antigenicity, instability index, immunogenicity and Ramachandran plot score of best binders of class I MHC.

### 3.6 Structural prediction and validation of epitopes

The three dimensional configuration of the predicted epitopes were produced by PEP-FOLD server. The stereo chemical features of the modelled epitopes were estimated by utilizing RAMPAGE server (**Fig. 2**). Cumulative distribution of Chi 1,2,3,4 and Phi-Psi torsional angles of the predicted peptides are qualitative indictors based on the percentile score of each residual conformation. Details regarding predictions obtained from RAMPAGE server are listed in **Table 2**. AGDSGFAAY, FVLAAVYRI, LSYFIASFR, and GFAAYSRYR exhibited good scores (100% in most favoured regions) and hence they are considered for furtherdocking analysis.

**Fig. 2.**
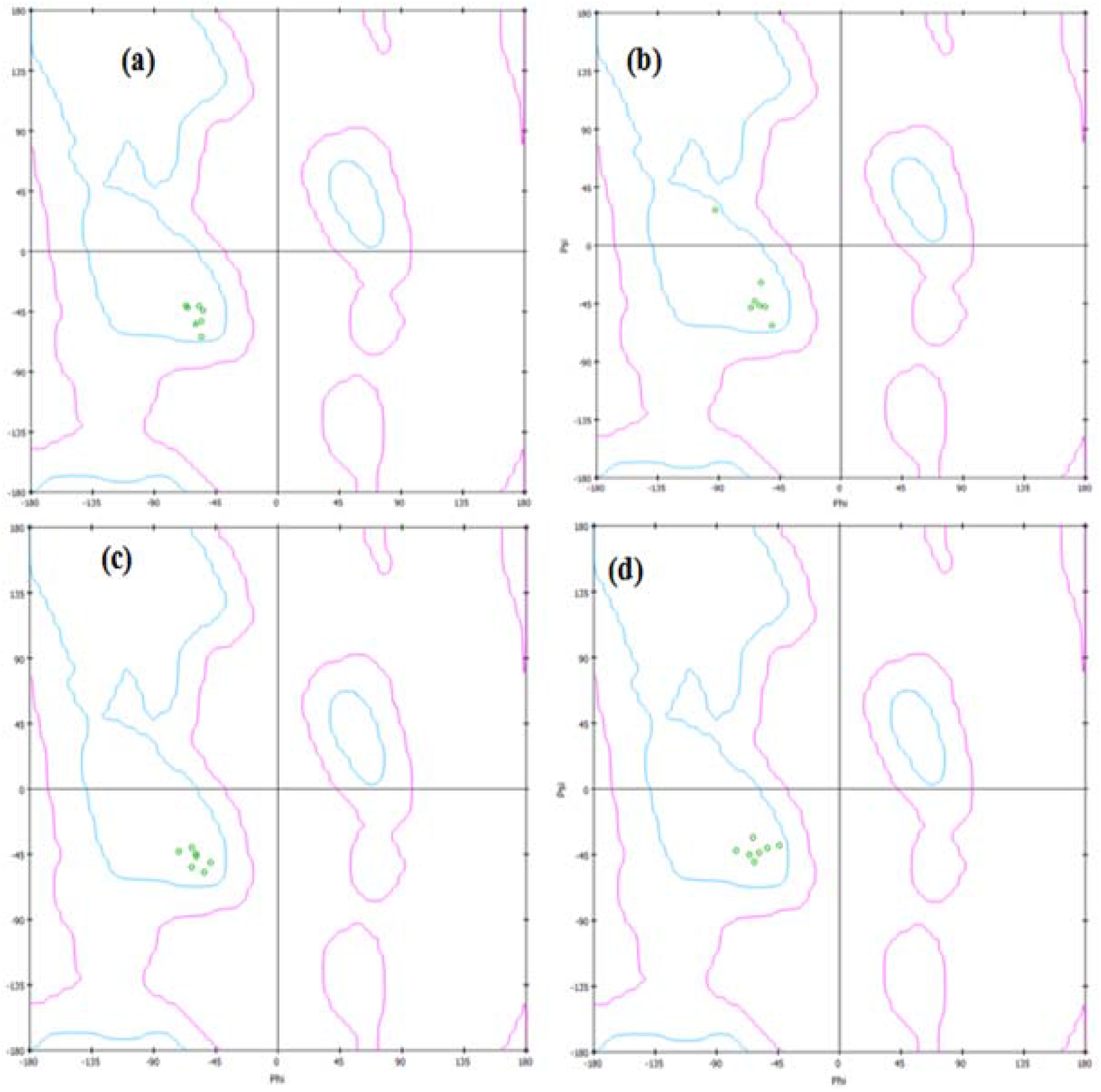
Validation of modelled structures using RAMPAGE (a) AGDSGFAAY (b) FVLAAVYRI(c) LSYFIASFR (d) GFAAYSRYR. The Ramachandran plot scores for these peptides were found to be 100%, which implies the stereochemical quality of the predicted peptides.

### 3.7 Molecular docking

The binding affinity of the predicted epitopes with their respective HLA-A (PDB ID: 1B0R) and HLA-B (PDB ID: 3C9N) were estimated using ClusPro server. Among, the four predicted epitope, FVLAAVYRI exhibited good binding energy (**Table 3**). The hydrogen bonding and electrostatic interactions of the peptide FVLAAVYRI were analyzed. The predicted peptide exhibited hydrogen bond interactions with the amino acid residues TRP147, PHE1, ALA4, GLN155 TYR7, and TYR159 which are significantly important for maximizing the stability of protein-ligand interactions (**Supplementary Table**). **Fig. 3** illustrates the 3D representation binding interactions of the epitope FVLAAVYRI with HLA-A and HLA-B of class I MHC molecules. The binding energies for four epitope are given the Table 3.

**Table 3:**
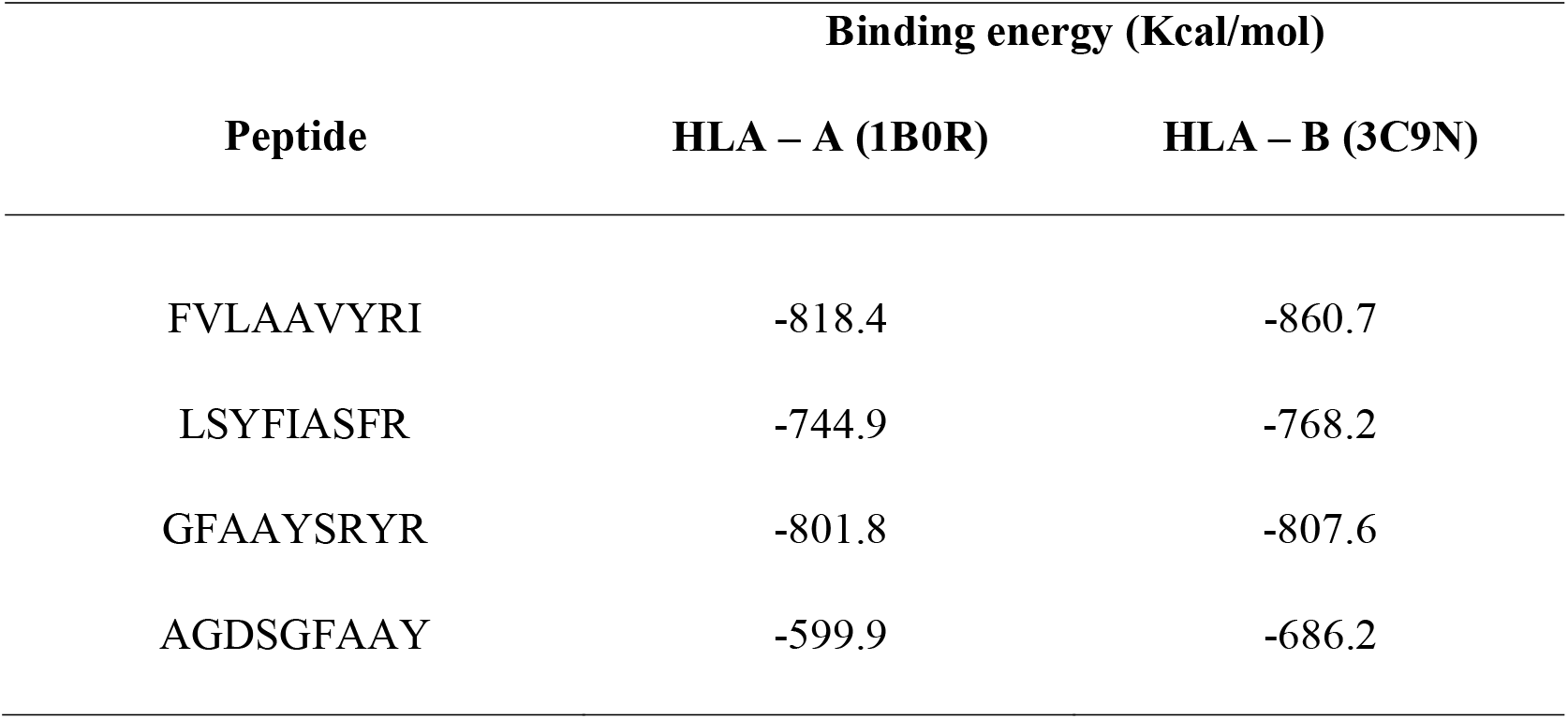
Binding energy values of epitopes

**Fig. 3.**
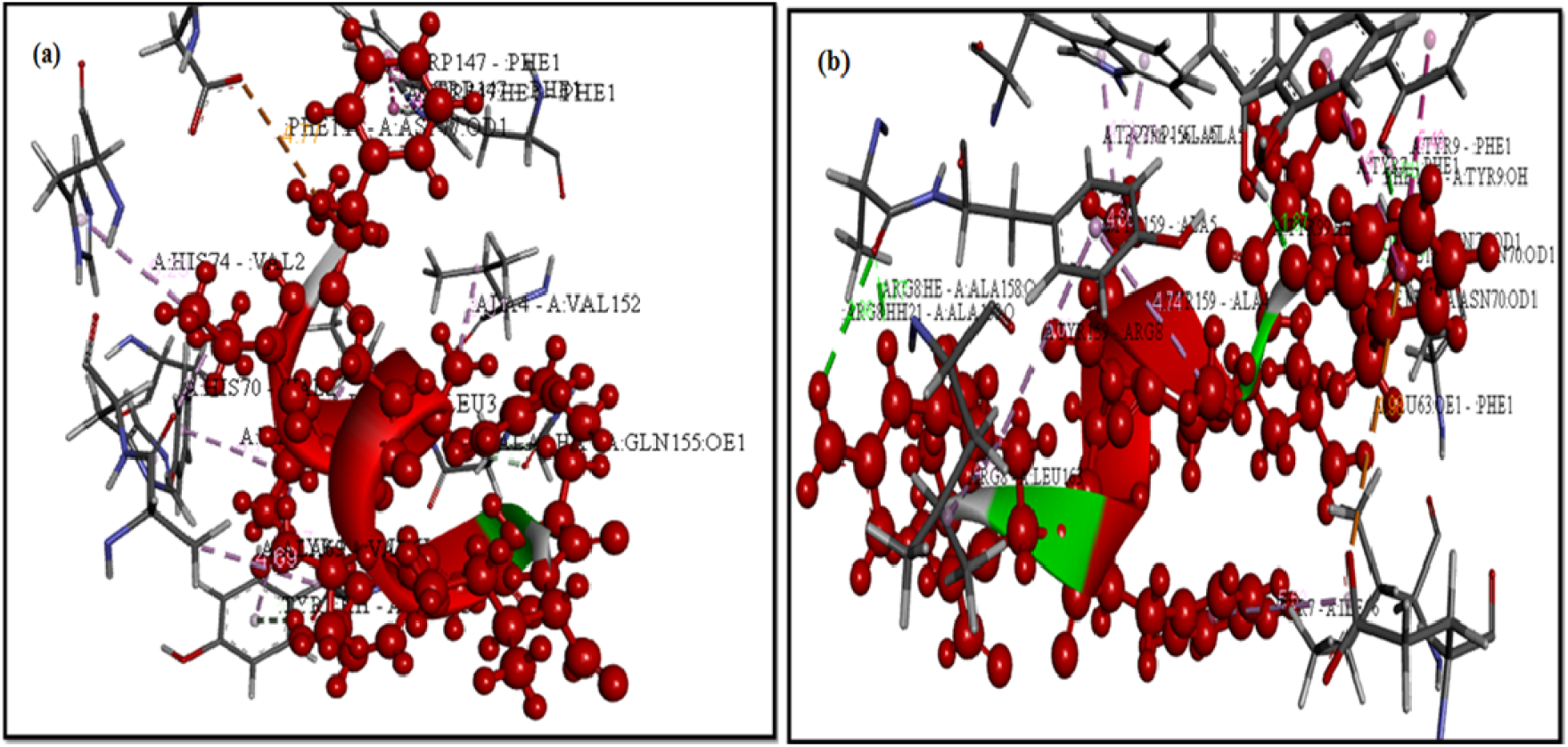
Binding interactions of FVLAAVYRI with (a) HLA-A and (b) HLA-B. Red colour ball and stick model indicates structural representation (solid model) of peptide FVLAAVYRI and key residues with HLA-A and HLA-B where Grey stick model indicates the amino acid residues, Green colour dotted lines represents hydrogen bonds; Purple and Pink colour dotted lines represent hydrophobic interactions; Orange colour dotted lines represents electrostatic interactions.

### 3.8 B-cell epitope prediction

B-cell epitope prediction is crucial to peptide driven vaccine design. The recognition of epitopes by B cells is based on features such as hydrophilicity, surface accessibility, flexibility, linearity and presence of beta turns. Bepipred tool aids in the determination of single scale amino acid propensity profiles B cell epitope is determined by the highest peak yellow region (threshold value >0.350) which corresponds to beta turns of the epitope (**Fig. 4a**). The beta turn commonly represents the surface accessible and hydrophilic regions of the proteins and epitopic regions predicted with the threshold value greater than 0.913. (**Fig. 4b**). The surface accessibility, hydrophilicity and antigenicity of predicted B-cell epitopes are shown **Fig.4c** and **Fig.4d** respectively. The antigenicity with the threshold value is about >1.0 and >0.970 thereby the predicted antigenic peptides are hydrophilic in nature. The amino acids and their action on identified B cell epitopes were determined with Kolasker and Tongaonkar tool (**Fig.4e**). It predicts the hydrophobic residues that are antigenic determined by physicochemical properties of amino acid residues with threshold value greater than 1.052. The Karplus & Schulz Flexibility prediction aided in the identification of flexible regions of predicted epitopes are also represented (**Fig. 4f**). The determined antigenic epitopes (threshold value >5.48) predicted based on the known temperature B-factors and also mobility of protein segments.

**Fig. 4.**
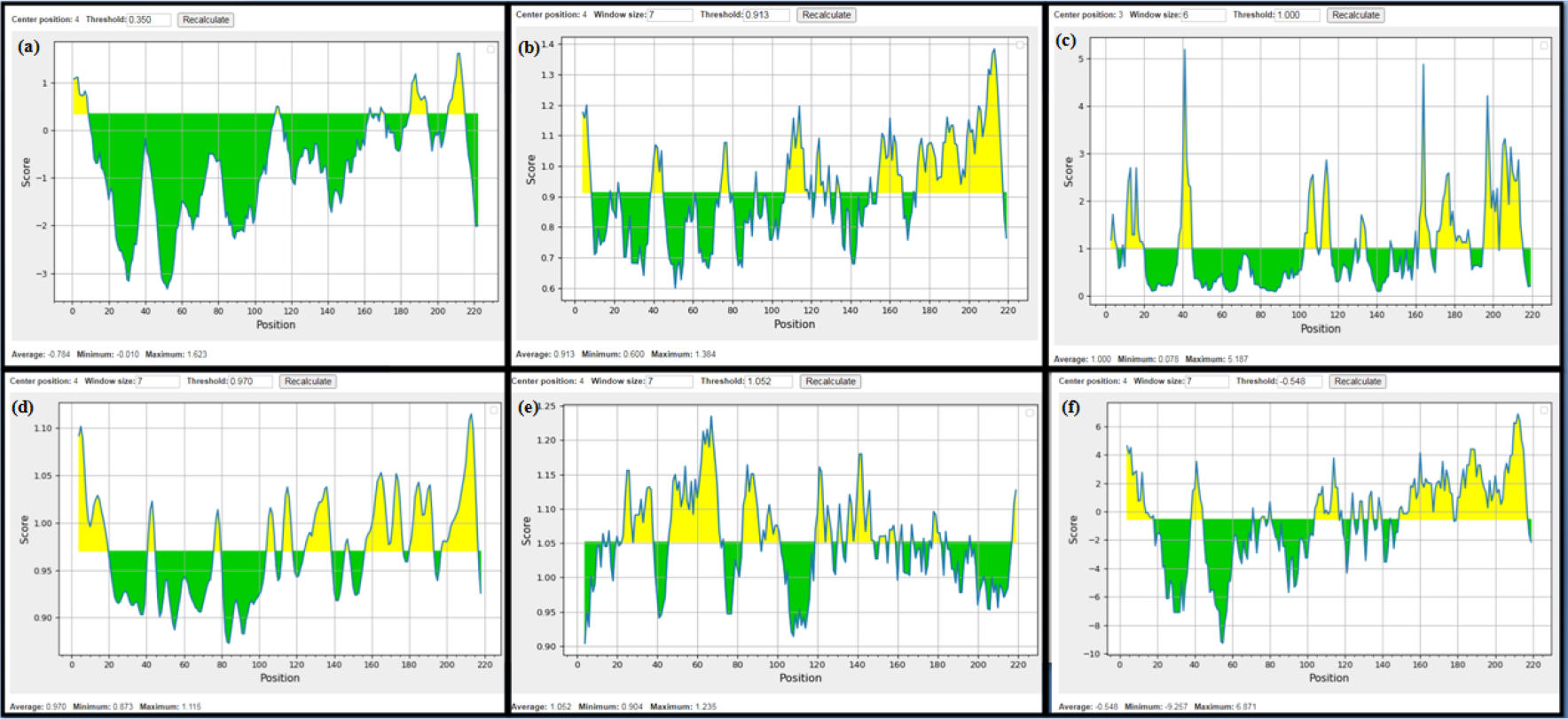
Membrane glycoprotein prediction using **(a)** Bepipred linear epitope with threshold value > 0.350 (**b)** Chou and Fasman beta-turn epitope with threshold value > 0.913 (**c)** Emini surface accessibility epitope with threshold value > 1.0 (**d)** Karplus & Schulz flexibilityepitope with threshold value > 0.950 (**e)** Kolaskar and Tongaonkar antigenicity epitope with threshold value > 1.052 and (**f)** Parker hydrophilicity epitope with threshold value > 0.548. The X-axis and Y-axis denote the sequence position and antigenic propensity respectively. The regions above the threshold value are antigenic, shown in yellow.

### 3.9 Molecular Dynamics study

The stability of docked peptide FVLAAVYRI with complex HLA- A and HLA -B molecules was further investigated by molecular dynamics simulations. The physicochemical properties were measured in the presence of water environment for 100 ns between 0 to 1000 frames. We observed the simulations of molecules are stable in the water environment with proper torsional periodicity. Molecular interaction of HLA-A between side chains of ligand peptide and protein surface residue forms instable configuration with the difference in the deviation distance from native position between 1.5A to 4.7 A at 45 ns time duration whereas the HLA-B shows stable configuration range of interaction with respect to time and distances of 0.74x higher than HLA-A complex. The system observes for the first half phase of the simulation periods are partially instable between the solvent and complex, complex and the ligand of HLA-A and HLA-B molecules. In the second half (50-100 ns) of the simulation periods shows that factors stable configuration. The dihedral angles between ligand and residual side chains complexes revealed conserved during 50ns simulation were 0.1 shown (**Supplementary Figure**) for HLA-A and HLA-B respectively. Thus this result proved that the stability of complex during simulation.

## 4.0 Discussion

Designing a new novel *insilico v*accine is a key to protect the global populations from the pandemic SARS-CoVID disease. Epitope/peptide vaccines are produced to induce immunity specific to pathogens by selective stimulation of antigen for B and T cells (Purcell et al. 2007). In this study, we propose to develop an epitope vaccine targeting membrane glycoprotein of SARS – CoV2, which plays an important role in viral assembly. Antigenicity, allergenicity as well as structural features of membrane protein was analysed by utilizing computational resources. The membrane protein possessed good antigenicity and least allergenicity. Further, T cell epitopes which could bind with class I MHC were predicted by exploiting IEDB tools. T cells immune response is considered to be more persistent than B cells, where the antibody memory response can be easily evaded by the antigen (Black et al. 2010). To efficiently protect against the invading pathogens, vaccines that produce cell-mediated responses are required. Also, CD 4+ and CD8+ T-cell responses play an important role in antiviral immunity(Sesardic 1993). The best binders of class I MHC were identified by considering their percentile score. Epitopes which possessed percentile score closer to 1 were identified as best binders. Epitopes such as YFIASFRLF, ATSRTLSYY, KLLEQWNLV, LVIGAVILR, YANRNRFLY, RIAGHHLGR, LAAVYRINW, FAYANRNRF, VATSRTLSY, AGDSGFAAY, RLFARTRSM, FVLAAVYRI, AVILRGHLR, LSYFIASFR, MACLVGLMW, GTITVEELK and GFAAYSRYRwere identified as best binders of class I MHC. The antigenicity, immunogenicity and stability index of the above mentioned epitopes were predicted. Epitopes such as LAAVYRINW, AGDSGFAAY, FVLAAVYRI, LSYFIASFR, GTITVEELK, and GFAAYSRYR exhibited good antigenic scores and immunogenicity scores compared to other epitopes. These epitopes three dimensional structure were generated and validated utilizing computational resources. Out of the six epitopes, AGDSGFAAY, FVLAAVYRI, LSYFIASFR, and GFAAYSRYR exhibited good stereo chemical profiles. Hence, these epitopes were subjected for docking studies to elucidate the binding affinity as well as binding energy with HLA-A and HLA-B of class I MHC. Epitope FVLAAVYRI exhibited good binding energy values when compared to all other epitopes. Behavior of post interacted complexes of HLA-A and B have been observed to be stable in the water solvent environment at various simulation times and distances. It functionally brings the exponential form of H bond interaction between each residue of the peptide (ligand) and the protein residues in the second phase of the simulation period. The consistency of the complex deviation was good enough to commit the functions of immune response mechanism. Hence, we propose FVLAAVYRI as a potential vaccine candidate for combating SARS-CoV infection. This can be strongly recommended as promising epitope vaccine.

Similarly, the immunoinformatic approaches were developed to design vaccines against Dengue (Ali et al. 2017), Hendra virus(Kamthania et al. 2019), Malaria (Pandey et al. 2018), Nipah virus (Ojha et al. 2019), *Klebsiella pneumonia*(Dar et al. 2019)*Pseudomonas aeruginosa* (Solanki et al. 2019) and cancerous antigens(Chauhan et al. 2019). The stimulation of host’s immune cells is triggered by the epitopes of B cells and T cells in the vaccine which in turn activates the other immune cells through complex signalling and MD simulations on peptide binding in the MHC binding groove was reported [32]. MD simulation studies showed complex had minimum changes during simulation and this study suggest that the complex was stable due to high-affinity epitopes at the binding sites.

## Conclusion

The current global pandemic SARS-COVID 19 is a devastating illness and it should be overcome through vaccination and immunogenicity of the individuals. *Insilico* vaccine design will be an effective strategy in dealing with the present situation. Epitope /peptide based vaccines are easy to produce and very effective in nature. It also reduces the cost and time required in producing vaccine and increases the specificity. Epitope vaccine design studies were reported against MERS-CoV, SARS-CoV and SARS-CoV-2, using *insilico* and immunoinformatic approaches by using B and T cell epitopes. These, immunoinformatic approaches can produce strong immune response against the pandemicSARS-COVID19. Epitopes chosen in this study are the ideal candidates to design a multiepitope peptide vaccine due to its antiallergenic and antigenic property. Its stability is assured by molecular docking and simulation studies. Epitope-based peptide vaccines can be systematised due to its known peptide structure and it can be easily altered to get multiepitope or conjugated structure infection. The *insilico* analysis of membrane glycoprotein predicts the preliminary level of analysis of mechanism of immune-biology of epitopes and its binding interactions with HLA-A and HLA-B allele. FVLAAVYRI is identified as a potential vaccine candidate for combating SARS-CoV which could be further validated by experimental immunization studies. The designed vaccine needs to be clinically verified in both *invitro* and *in vivo* studies to assure its safety and to halt this pandemic viral disease from spreading even further.

## ACKNOWLEDGEMENTS

We appreciate the contributions of all those who participated in this research and the comments of the reviewers of this manuscript.

## COMPLIANCE WITH ETHICAL STANDARDS

### FUNDING

None

### CONFLICTS OF INTEREST

On behalf of all authors, the corresponding author states that there is no conflict of interest. Ethical approval: This article does not contain any studies with human participants performed by any of the authors.

This article does not contain any studies with animals performed by any of the authors.

This article does not contain any studies with human participants or animals performed by any of the authors.

