## Supplementary file for "Immunoinformatic approach to design a vaccine against SARS-COV-2 membrane glycoprotein"

**Supporting Information**

**Table 1S**: **MHC Class I alleles for SARS-CoV2 membrane glycoprotein protein**.

| **Start** | **End** | **Length** | **Peptide** | **Allele** | **Score** | **Rank** |
| --- | --- | --- | --- | --- | --- | --- |
| 95 | 103 | 9 | YFIASFRLF | HLA-A*23:01 | 0.974862 | 0.01 |
|  |  |  |  | HLA-A*24:02 | 0.967974 | 0.01 |
|  |  |  |  | HLA-A*32:01 | 0.14648 | 0.76 |
|  |  |  |  | HLA-A*26:01 | 0.119241 | 0.75 |
| 94 | 102 | 9 | SYFIASFRL | HLA-A*23:01 | 0.952022 | 0.03 |
|  |  |  |  | HLA-A*24:02 | 0.939447 | 0.03 |
|  |  |  |  | HLA-A*32:01 | 0.186478 | 0.58 |
| 171 | 179 | 9 | ATSRTLSYY | HLA-A*01:01 | 0.921616 | 0.06 |
|  |  |  |  | HLA-A*30:02 | 0.874028 | 0.01 |
|  |  |  |  | HLA-A*11:01 | 0.697222 | 0.15 |
|  |  |  |  | HLA-A*03:01 | 0.535041 | 0.29 |
|  |  |  |  | HLA-B*58:01 | 0.533599 | 0.31 |
|  |  |  |  | HLA-A*26:01 | 0.447843 | 0.16 |
|  |  |  |  | HLA-A*68:01 | 0.327433 | 0.84 |
|  |  |  |  | HLA-A*30:01 | 0.295806 | 0.37 |
|  |  |  |  | HLA-A*32:01 | 0.145151 | 0.77 |
| 15 | 23 | 9 | KLLEQWNLV | HLA-A*02:01 | 0.903776 | 0.04 |
|  |  |  |  | HLA-A*32:01 | 0.283692 | 0.37 |
| 138 | 146 | 9 | LVIGAVILR | HLA-A*68:01 | 0.901219 | 0.02 |
|  |  |  |  | HLA-A*11:01 | 0.597484 | 0.23 |
|  |  |  |  | HLA-A*31:01 | 0.570689 | 0.18 |
|  |  |  |  | HLA-A*33:01 | 0.497373 | 0.1 |
|  |  |  |  | HLA-A*03:01 | 0.357347 | 0.53 |
| 39 | 47 | 9 | YANRNRFLY | HLA-B*35:01 | 0.891748 | 0.04 |
|  |  |  |  | HLA-A*01:01 | 0.86633 | 0.1 |
|  |  |  |  | HLA-A*30:02 | 0.790439 | 0.03 |
|  |  |  |  | HLA-B*58:01 | 0.740428 | 0.14 |
|  |  |  |  | HLA-B*53:01 | 0.567363 | 0.12 |
|  |  |  |  | HLA-A*68:01 | 0.527198 | 0.38 |
|  |  |  |  | HLA-A*26:01 | 0.336561 | 0.23 |
| 150 | 158 | 9 | RIAGHHLGR | HLA-A*31:01 | 0.882665 | 0.01 |
|  |  |  |  | HLA-A*03:01 | 0.812219 | 0.05 |
|  |  |  |  | HLA-A*11:01 | 0.607438 | 0.23 |
|  |  |  |  | HLA-A*68:01 | 0.461478 | 0.51 |
|  |  |  |  | HLA-A*30:01 | 0.306423 | 0.34 |
|  |  |  |  | HLA-A*33:01 | 0.261419 | 0.39 |
| 188 | 196 | 9 | AGDSGFAAY | HLA-A*01:01 | 0.821914 | 0.13 |
|  |  |  |  | HLA-A*30:02 | 0.652631 | 0.1 |
|  |  |  |  | HLA-B*35:01 | 0.405673 | 0.34 |
| 67 | 75 | 9 | LAAVYRINW | HLA-B*58:01 | 0.871882 | 0.05 |
|  |  |  |  | HLA-B*53:01 | 0.49967 | 0.15 |
|  |  |  |  | HLA-A*32:01 | 0.143564 | 0.77 |
| 37 | 45 | 9 | FAYANRNRF | HLA-B*35:01 | 0.861293 | 0.05 |
|  |  |  |  | HLA-B*53:01 | 0.715917 | 0.05 |
|  |  |  |  | HLA-B*58:01 | 0.644898 | 0.2 |
|  |  |  |  | HLA-B*51:01 | 0.589489 | 0.14 |
|  |  |  |  | HLA-A*23:01 | 0.406378 | 0.48 |
|  |  |  |  | HLA-B*08:01 | 0.339201 | 0.41 |
|  |  |  |  | HLA-A*24:02 | 0.316854 | 0.7 |
| 170 | 178 | 9 | VATSRTLSY | HLA-B*35:01 | 0.85887 | 0.05 |
|  |  |  |  | HLA-A*30:02 | 0.682184 | 0.08 |
|  |  |  |  | HLA-A*01:01 | 0.582992 | 0.3 |
|  |  |  |  | HLA-B*58:01 | 0.517414 | 0.33 |
|  |  |  |  | HLA-B*53:01 | 0.352793 | 0.27 |
|  |  |  |  | HLA-A*26:01 | 0.265007 | 0.33 |
| 84 | 92 | 9 | MACLVGLMW | HLA-B*58:01 | 0.776528 | 0.11 |
|  |  |  |  | HLA-B*53:01 | 0.393241 | 0.23 |
| 142 | 150 | 9 | AVILRGHLR | HLA-A*31:01 | 0.790975 | 0.04 |
|  |  |  |  | HLA-A*68:01 | 0.64001 | 0.22 |
|  |  |  |  | HLA-A*33:01 | 0.47081 | 0.12 |
|  |  |  |  | HLA-A*11:01 | 0.457745 | 0.38 |
|  |  |  |  | HLA-A*03:01 | 0.289192 | 0.67 |
|  |  |  |  | HLA-A*30:01 | 0.172517 | 0.97 |
| 101 | 109 | 9 | RLFARTRSM | HLA-B*08:01 | 0.799697 | 0.03 |
|  |  |  |  | HLA-A*32:01 | 0.771969 | 0.03 |
|  |  |  |  | HLA-A*30:01 | 0.41936 | 0.14 |
|  |  |  |  | HLA-B*07:02 | 0.296987 | 0.45 |
|  |  |  |  | HLA-A*02:01 | 0.22354 | 0.98 |
|  |  |  |  | HLA-A*03:01 | 0.218454 | 0.87 |
|  |  |  |  | HLA-A*30:02 | 0.20334 | 0.78 |
| 65 | 73 | 9 | FVLAAVYRI | HLA-A*02:01 | 0.674241 | 0.2 |
|  |  |  |  | HLA-A*68:02 | 0.531024 | 0.21 |
|  |  |  |  | HLA-B*51:01 | 0.26847 | 0.58 |
|  |  |  |  | HLA-A*32:01 | 0.219935 | 0.49 |
| 93 | 101 | 9 | LSYFIASFR | HLA-A*68:01 | 0.78421 | 0.08 |
|  |  |  |  | HLA-A*31:01 | 0.625748 | 0.13 |
|  |  |  |  | HLA-A*11:01 | 0.472446 | 0.36 |
|  |  |  |  | HLA-A*33:01 | 0.471942 | 0.12 |
|  |  |  |  | HLA-A*03:01 | 0.270028 | 0.71 |
| 6 | 14 | 9 | GTITVEELK | HLA-A*11:01 | 0.769234 | 0.09 |
|  |  |  |  | HLA-A*68:01 | 0.645005 | 0.22 |
|  |  |  |  | HLA-A*03:01 | 0.314923 | 0.62 |
| 192 | 200 | 9 | GFAAYSRYR | HLA-A*31:01 | 0.67837 | 0.09 |
|  |  |  |  | HLA-A*33:01 | 0.33929 | 0.25 |

**Table 2S : Electrostatic interactions of best docked complex**

| **Interactions** | **HLA -A** | | **HLA-B** | |
| --- | --- | --- | --- | --- |
|  | **Donor ⋅⋅⋅Acceptor** | **Distance (Ǻ)** | **Donor ⋅⋅⋅Acceptor** | **Distance (Ǻ)** |
| **Electrostatic**  **/Pi-cation** | PHE1(N)⋅⋅⋅ASP77(OD1) | 4.77 | GLU63(OE1) ⋅⋅⋅PHE1 | 4.98 |
| **Hydrogen**  **interaction** | TRP147(HE1) ⋅⋅⋅PHE1  ALA4(HA) ⋅⋅⋅GLN155(OE1)  TYR7[22]⋅⋅⋅TYR159 | 2.16  2.62  3.07 | ARG8(HH21) ⋅⋅⋅ALA158(O)  ARG8(HE) ⋅⋅⋅ALA158(O)  TYR99[22]⋅⋅⋅PHE1(O)  PHE1(H2) ⋅⋅⋅TYR(OH)  VAL2(H) ⋅⋅⋅ASN70(OD1)  PHE1(H1) ⋅⋅⋅ASN70(OD1)  PHE1(HA) ⋅⋅⋅ASN70(OD1) | 2.98  1.77  1.87  1.80  1.73  1.83  2.78 |
| **Hydrophobic interaction** | | | | |
| Pi-pi –T-shaped interaction | TRP147(HE1)⋅⋅⋅PHE1  TRP147⋅⋅⋅PHE1 | 4.31  5.41 | TYR9⋅⋅⋅PHE1  TYR7⋅⋅⋅PHE1 | 5.49  4.72 |
| Alkyl and pi-alkyl | HIS74⋅⋅⋅VAL2  HIS70⋅⋅⋅VAL2  LEU156⋅⋅⋅LEU3  ALA4⋅⋅⋅VAL152  TYR99⋅⋅⋅LEU3  TYR159⋅⋅⋅LEU3  ALA69⋅⋅⋅VAL6 | 5.29  3.96  5.09  3.81  4.71  5.37  4.69 | TYR7⋅⋅⋅ILE66  ARG8⋅⋅⋅LEU163  TYR159⋅⋅⋅ARG8  TYR159⋅⋅⋅ALA4  TYR159⋅⋅⋅ALA4  TRP156⋅⋅⋅ALA5  TRP156⋅⋅⋅ALA5 | 5.50  4.95  5.08  4.74  4.66  3.43  4.01 |

**FIGURE**

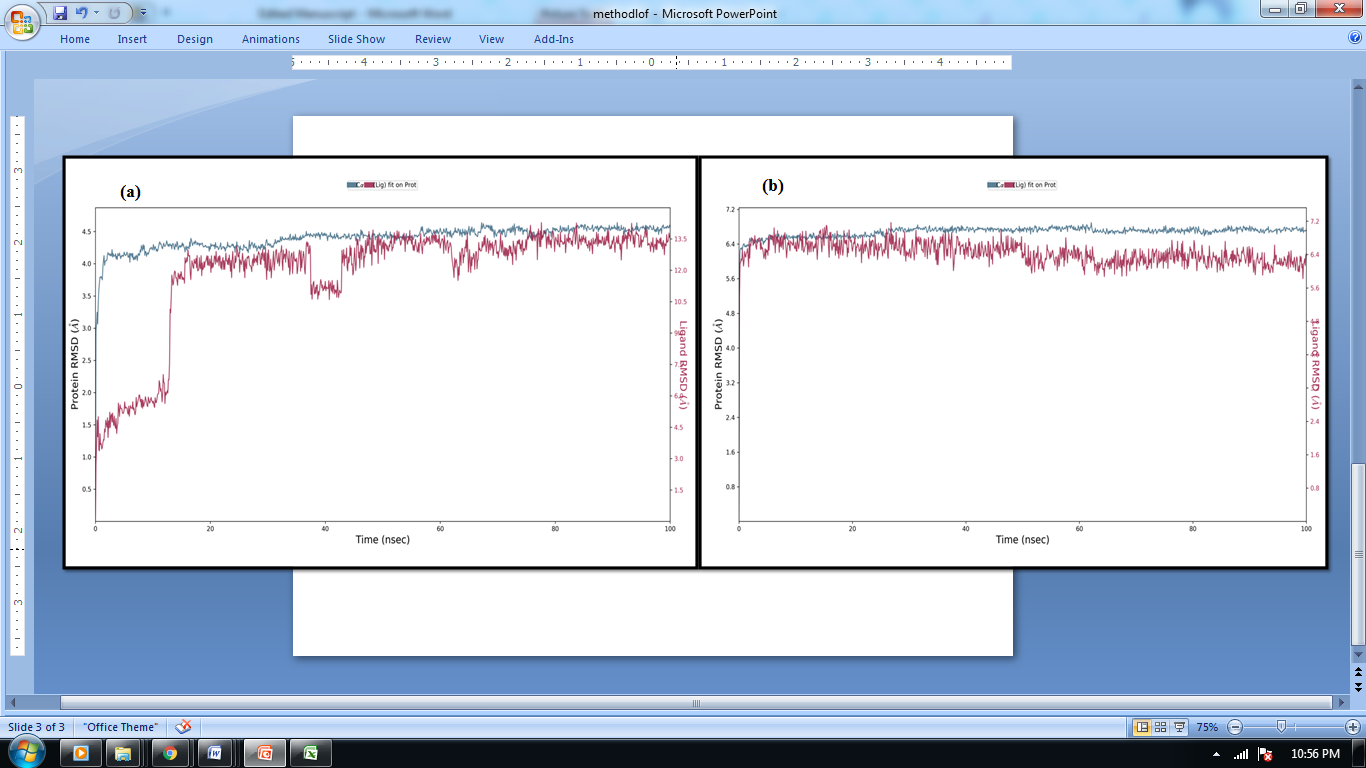

**Fig 1S:** MD simulation of **(a)** HLA-A and **(b)** HLA -B complexed with 9mer peptide – FVLAAVYRI. Deviation of complete complex and fluctuation of individual residues were calculated from RMSD and RMSF respectively. The simulation time of the complex was calculated from the conformation of alpha carbon position of each residue.
